# Structural Basis for the Assembly of Amyloid Fibrils by the Master Cell-Signaling Regulator Human Receptor-Interacting Protein Kinase 1

**DOI:** 10.1101/2025.01.13.630173

**Authors:** Paula Polonio, Jorge Pedro López-Alonso, Hanxing Jiang, Fátima Escobedo, Gustavo Titaux, Iban Ubarretxena-Belandia, Miguel Mompeán

**Affiliations:** Instituto de Química Física Blas-Cabrera (IQF-CSIC), Madrid, Spain; Instituto Biofisika (UPV/EHU, CSIC), Leioa, Spain; Basque Resource for Electron Microscopy, Leioa, Spain; Ikerbasque Foundation for Science, Bilbao, Spain

## Abstract

Amyloid fibrils, typically associated with neurodegenerative diseases, also play critical roles as functional assemblies in biological processes. The RIP homotypic interaction motifs (RHIMs) in receptor-interacting protein kinases 1 and 3 (RIPK1 and RIPK3) are essential for necroptosis, orchestrating the formation of amyloid-like fibrils that assemble into necrosomes. These supramolecular complexes propagate cell death signals and activate effectors like MLKL. While the structures of human RIPK3 (hRIPK3) homomeric fibrils and RIPK1-RIPK3 heteromeric fibrils have been resolved, the atomic structure of human RIPK1 (hRIPK1) homomeric fibrils has remained elusive.

Here, we present a high-resolution structure of hRIPK1 RHIM-mediated amyloid fibrils, determined using an integrative approach combining cryoprobe-detected solid-state nuclear magnetic resonance spectroscopy and cryo-electron microscopy. The fibrils adopt an N-shaped amyloid fold consisting of three β-sheets stabilized by the conserved IQIG RHIM motif through hydrophobic interactions and hydrogen bonding. A key hydrogen bond between N545 and G542 closes the β2-β3 loop, resulting in denser side-chain packing compared to hRIPK3 homomeric fibrils. This structural feature likely contributes to the compact architecture of hRIPK1 fibrils, in contrast to the more relaxed S-shaped fold observed in hRIPK3.

These findings provide structural insights into how hRIPK1 homomeric fibrils nucleate hRIPK3 recruitment and fibrillization during necroptosis, offering broader perspectives on the molecular principles governing RHIM-mediated amyloid assembly and functional amyloids.

## Introduction

Receptor-Interacting Protein Kinase 1 (RIPK1) is a master regulator of cell signaling, orchestrating diverse cellular outcomes depending on the context. Through polyubiquitination by E3 ligases, RIPK1 promotes cell proliferation and differentiation (1). In contrast, when polyubiquitination is inhibited, RIPK1 forms a secondary cytosolic complex with FADD and caspase-8 to trigger apoptosis (2,3). Furthermore, under conditions where caspase activity is blocked – such as during viral infections – RIPK1 interacts with RIPK3 to trigger necroptosis (4,5). These versatile functions rely on the multidomain architecture of RIPK1, which comprises a N-terminal kinase domain, an intermediate disordered region harboring a RIP Homotypic Interaction Motif (RHIM), and a C-terminal Death Domain (**Fig. 1a**). The RHIM motif, which is a tetrapeptide with consensus sequence (V/I)-Q-(V/I/L/C)-G, is essential to signal necroptosis.

**Fig. 1.**
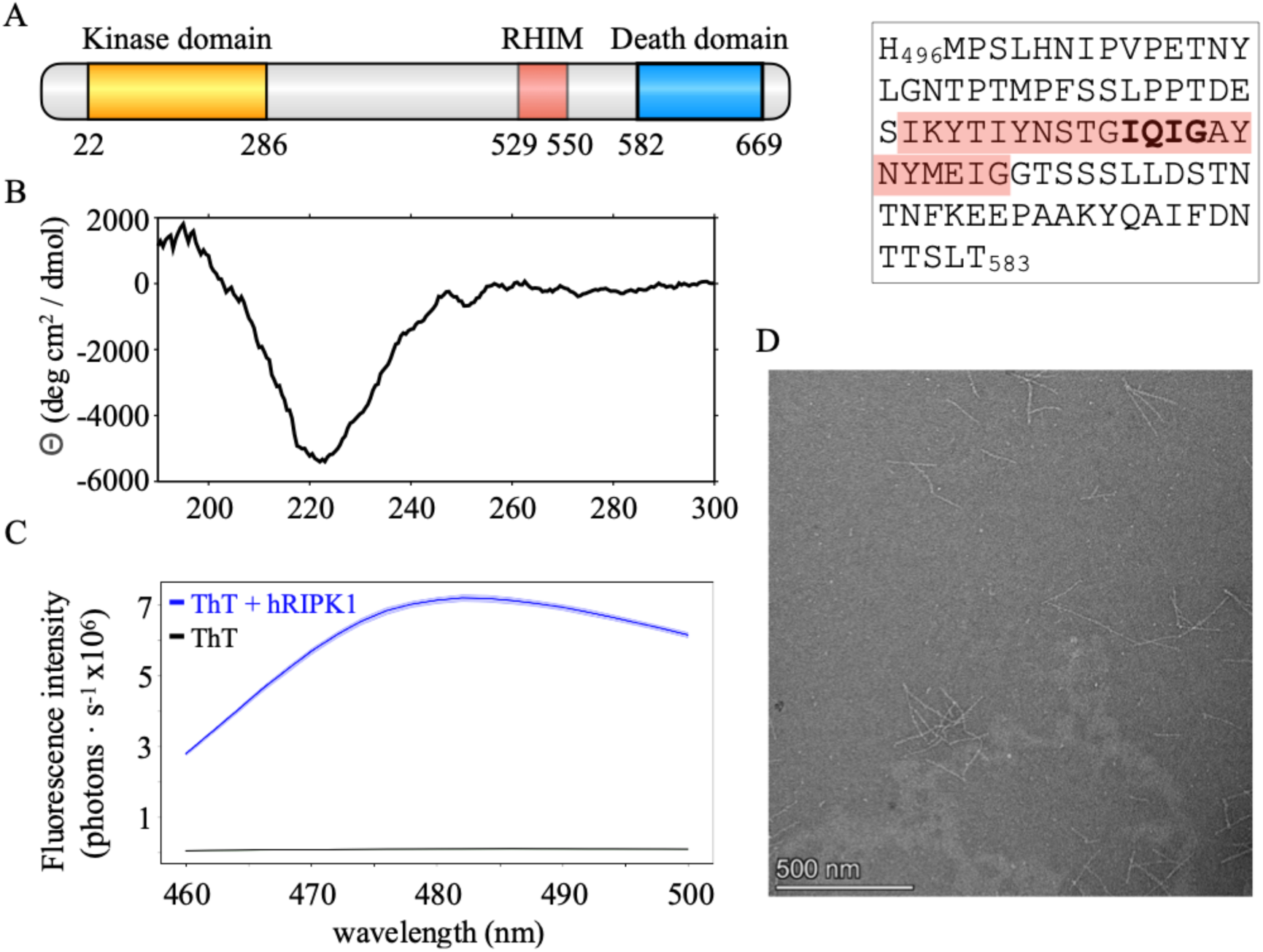
RHIM hRIPK1 forms amyloid fibrils. **a** Schematic of the primary structure of hRIPK1 depicting the N-terminal kinase domain, the intermediate RHIM-harboring domain, and the C-terminal Death Domain. In this work we focused on the intermediate RHIM region of hRIPK1 encompassing amino acids 496-583. The amino acid sequence of RHIM hRIPK1 is shown on the right with those residues that have been shown to assemble into heteromeric RIPK1-RIPK3 fibrils highlighted in pink, and with the RHIM IQIG sequence in bold. **b** Room temperature CD spectrum of RHIM hRIPK1 in water at pH 7.4 displaying a pronounced minimum at ∼220 nm, indicative of high β-strand secondary structure content. **c** Room temperature ThT fluorescence emission spectrum in water at pH 7.4 in the absence (black) and presence (blue) of RHIM hRIPK1. Increased fluorescence by ThT is a reporter of RHIM hRIPK1 amyloid formation. **d** Negative-stain TEM imaging reveals RHIM hRIPK1 unbranched fibrils. Scale bar is shown.

Necroptosis is a programmed cell death pathway that plays a crucial role in maintaining tissue homeostasis, defending against pathogens, and resolving inflammation (6–9). Unlike apoptosis, which involves caspase-mediated signaling, necroptosis is a regulated form of necrotic cell death that relies on functional amyloid fibril assemblies and is dependent on the kinase activities of RIPK1 and RIPK3 (4,10). These RIPK kinases assemble into functional amyloids via their RHIM motifs (11,12), forming a heteromeric (RIPK1-RIPK3) fibrillar complex known as the canonical necrosome. This necrosome acts as a signaling platform that templates and seeds the assembly of RIPK3 homomeric fibrils, which in turn recruit and oligomerize the mixed lineage kinase domain-like protein (MLKL). MLKL oligomers then separate from RIPK3 and translocate to the plasma membrane, disrupting its integrity and ultimately inducing cell death (13–15).

Activation of MLKL for necroptosis execution can also occur through non-canonical necrosomes, which are formed by the heteromeric assembly of RIPK3 with other proteins distinct from RIPK1 that also contain RHIM motifs, such as Z-DNA Binding Protein 1 (ZBP1) (16). Despite their distinct composition, both canonical (RIPK1-RIPK3) and non-canonical (e.g. ZBP1-RIPK3) necrosomes converge on the activation of homomeric RIPK3 fibrils and subsequent oligomerization of MLKL for membrane rupture (16).

While the canonical RIPK1-RIPK3 and non-canonical ZBP1-RIPK3 necrosomes appear to function analogously (i.e. templating downstream homomeric assembly of RIPK3 fibrils to activate MLKL), recent studies have demonstrated that this is not actually the case. Particularly, it has been shown that, in human cells, the RHIM-mediated association of hZBP1 and hRIPK3 critically depends on the prior assembly of a stable hRIPK1-hZBP1 complex, which also involves the RHIM motif of hRIPK1 (17). Interestingly, murine systems bypass this dependency, highlighting differences in necroptotic signaling across species (17).

Beyond necroptosis, the RHIM of hRIPK1 has been also recently shown to be essential for hRIPK1-hZBP1 interaction during necroptosis-independent inflammation (18), further underscoring the broader regulatory significance of hRIPK1 and its RHIM-driven assemblies. The promiscuity of hRIPK1 as “an essential signaling node” (19) is underpinned by its ability to form functional amyloid fibrils via its RHIM motif. These fibrils act as tightly regulated scaffolds that facilitate cell signaling, distinguishing them from the pathological amyloids associated with neurodegenerative disorders (20,21).

By serving as reversible platforms for the assembly of heteromeric and homomeric complexes, the RHIM of RIPK1 enables both canonical and non-canonical necrosome signaling, and it also supports necroptosis independent pathways (18). Structural studies have provided insights into RHIM-driven fibrils, revealing an S-shaped fold for hRIPK3 (22) and an N-shaped fold for mouse RIPK1 (mRIPK1) (23). In both mRIPK1 and hRIPK3 homomeric fibrils, the RHIM tetrapeptides (IQIG in RIPK1 and VQVG in RIPK3) adopt similar conformations to those observed in the human RIPK1-RIPK3 heteromeric complex (12). However, the structure of hRIPK1 RHIM-driven homomeric fibrils remains unknown, limiting our understanding of how hRIPK1 initiates and regulates the reversible assembly of both homomeric fibrils and the recruitment of hRIPK3 and hZBP1 into heteromeric amyloids to control critical necroptotic and inflammatory signaling pathways in humans.

We present here a high-resolution structure of hRIPK1 RHIM fibrils determined by a combination of CPMAS cryoprobe-detected solid-state nuclear magnetic resonance (SSNMR) spectroscopy and cryo-electron microscopy (cryo-EM). The structure unveils the structural basis for the assembly of compact N-shaped hRIPK1 amyloid fibrils providing insight into the molecular mechanisms of RHIM-mediated signaling and a rational for hRIPK3 activation that advances our understanding of necrosome assembly.

## Results

### Formation of RHIM hRIPK1 fibrils

To determine the structure of hRIPK1 fibrils, we produced a fragment of hRIPK1 spanning residues 496-583 that includes the RHIM region (**Fig. 1a**). This fragment has consistently been shown to act as a *bona fide* assembling domain in a number of studies on homo- and heteromeric RHIM-driven interactions (11,12,24,25). Dialysis of the hRIPK1 fragment against Tris-HCl buffer (pH 7.4) at room temperature to remove the SDS employed during purification led to turbidity. This turbidity suggested fibril formation. Circular dichroism (CD) spectroscopy of the turbid material revealed a pronounced minimum at ∼220 nm (**Fig. 1b**), indicative of extensive β-sheet secondary structure content (26). Thioflavin T (ThT) binding assays were consistent with the formation of amyloid-like fibrils by hRIPK1 (496–583) (27) (**Fig. 1c**), which were shown to be homogeneous in width and length by negative-stain transmission electron microscopy (**Fig. 1d**).

### SSNMR reveals the extended β-strand conformation of the fibril core

SSNMR is well-suited for structure determination of hydrated amyloid fibrils. This technique traditionally relies on significant amounts of different isotopically labelled samples to distinguish intra- and/from intermolecular interactions (28). Here we avoided laborious sample preparation schemes by exploiting the enhanced sensitivity afforded by a CPMAS HCN cryoprobe specifically designed for biosolids (29), and were able to record all SSNMR spectra from a single isotopically diluted hRIPK1 fibril sample at a 1:4 molar ratio of ^13^C,^15^N-labeled to unlabeled protein. The labeled and unlabeled proteins were mixed under denaturing conditions and dialyzed to induce fibril formation. This strategy ensured that, statistically, one in every four stacked monomers along the fibril axis is isotopically labeled (and thus NMR-visible), effectively minimizing intermolecular contributions and enabling characterization of the protomeric conformation within the fibril.

A minimum set of three 3D SSNMR experiments enabled sequential backbone assignments with confidence. In particular, NCACX, NCOCX, and CANCOCX experiments provided a detailed map of backbone and side-chain resonances (**Fig. 2a**). Due to the isotopic dilution, all signals can be regarded as intramolecular and thus arising from atomic nuclei in one protomer within the fibril. This level of data collection would be impractical with conventional CPMAS probes unless fully isotopically labeled samples were used, which could in turn reintroduce intermolecular ambiguities (e.g. a CA-CB contact for a given residue could be either intramolecular or intermolecular due to monomer stacking within the fibril).

**Fig. 2.**
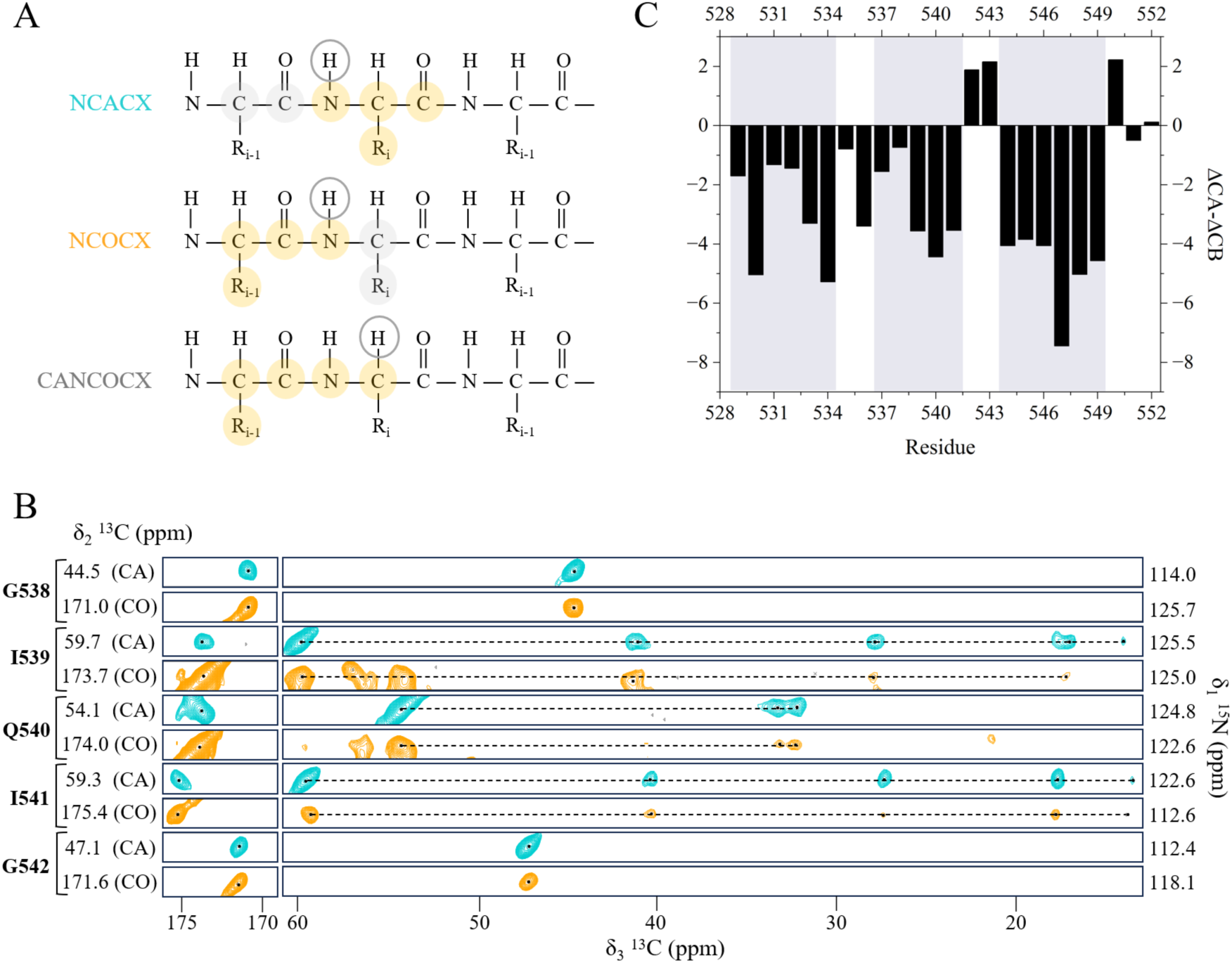
The extended β-strand conformation of the RHIM hRIPK1 fibril core. **a** Schematic representation of the magnetization transfer in the three 3D experiments used for backbone sequential assignment. **b** Strip plots showing the sequential connectivity within the core of RIPK1 RHIM (GIQIG), extracted from NCACX (blue) and NCOCX (orange) experiments at 14.1 T, 298 K and MAS rate of 14kHz. **c** Secondary Structure Propensity (SSP) for the hRIPK1 amyloid spanning residues I529-T552 highlighting three β-strands (gray shadow) encompassing residues I529-Y534 (β1), T537-I541 (β2), and Y544-I549 (β3), with residues N535-S536 adopting a kinked conformation between β1 and β2, and G542-A543 forming a turn between β2 and β3.

The NCACX experiment yielded correlations within each residue, capturing N*_i_*, CA*_i_*, and CX*_i_* signals for the residue *i*, where CX*_i_* represents both CO*_i_* and ^13^C*_i_* side-chain atoms. The NCOCX experiment linked the ^13^C atoms of a residue *i* to the nitrogen atom of the next residue (N*_i+1_*), while the CANCOCX experiment bridged CA*_i+1_* and N*_i+1_* with the ^13^C atoms of residue *i*, enabling sequential connectivity throughout the fibril core (**Fig. 2a,b**). Using this approach, we obtained site specific chemical shift assignments for the stretch spanning residues 529-552 that encompasses the I539-Q540-I541-G542 RHIM motif (**Fig. 1a**). Notably, since these experiments rely on dipolar interactions for magnetization transfer, they selectively detect residues in the rigid amyloid core, which remains observable even under hydrated and physiological temperature conditions – where more flexible regions typically experience increased dynamics and become undetectable.

To interrogate the conformations adopted by the hRIPK1 I529-T552 fibril core under these conditions, we next analyzed the difference between CA and CB experimental chemical shifts (CA*_exp_* and CB*_exp_*) with their predicted values for a random coil sequence (CA*_rc_* and CB*_rc_*), that is: Δ(CA*_exp_* – CA*_rc_*) – Δ(CB*_exp_* – CB*_rc_*). These values provide a reliable indicator of secondary structure propensities, or SSP (30), and consistently negative SSP values that are indicative of extended β-strand conformations were observed for most residues within the core region of the fibril (**Fig. 2c**).

### SSNMR structure of RHIM hRIPK1 fibril protomers

Having delineated the amyloid core of hRIPK1, we next sought to construct a detailed structural model the fibril protomers using 2D ^13^C-^13^C CORD experiments recorded with 20, 100, and 500 ms mixing times. At short mixing times (20 ms), these experiments primarily exhibited intra-residue correlations, such as CA–CB or CA–CO within the same residue. Intermediate mixing times (100 ms) revealed sequential correlations between consecutive residues (*i* and *i+1*), confirming the fibril’s backbone connectivity and providing further assignments, such as aromatic resonances from Tyr residues. At long mixing times (500 ms), correlations between residues separated by two, three or more positions in the sequence were detected, providing long-range distance constraints that are essential for defining the overall fold of hRIPK1 monomers within the fibril core. For example, strong correlations were observed between G542 CA and N545 CA, as well as for G538 CA and M547 CG, highlighting their proximity within the fibril core (**Fig. 3a**).

**Fig. 3.**
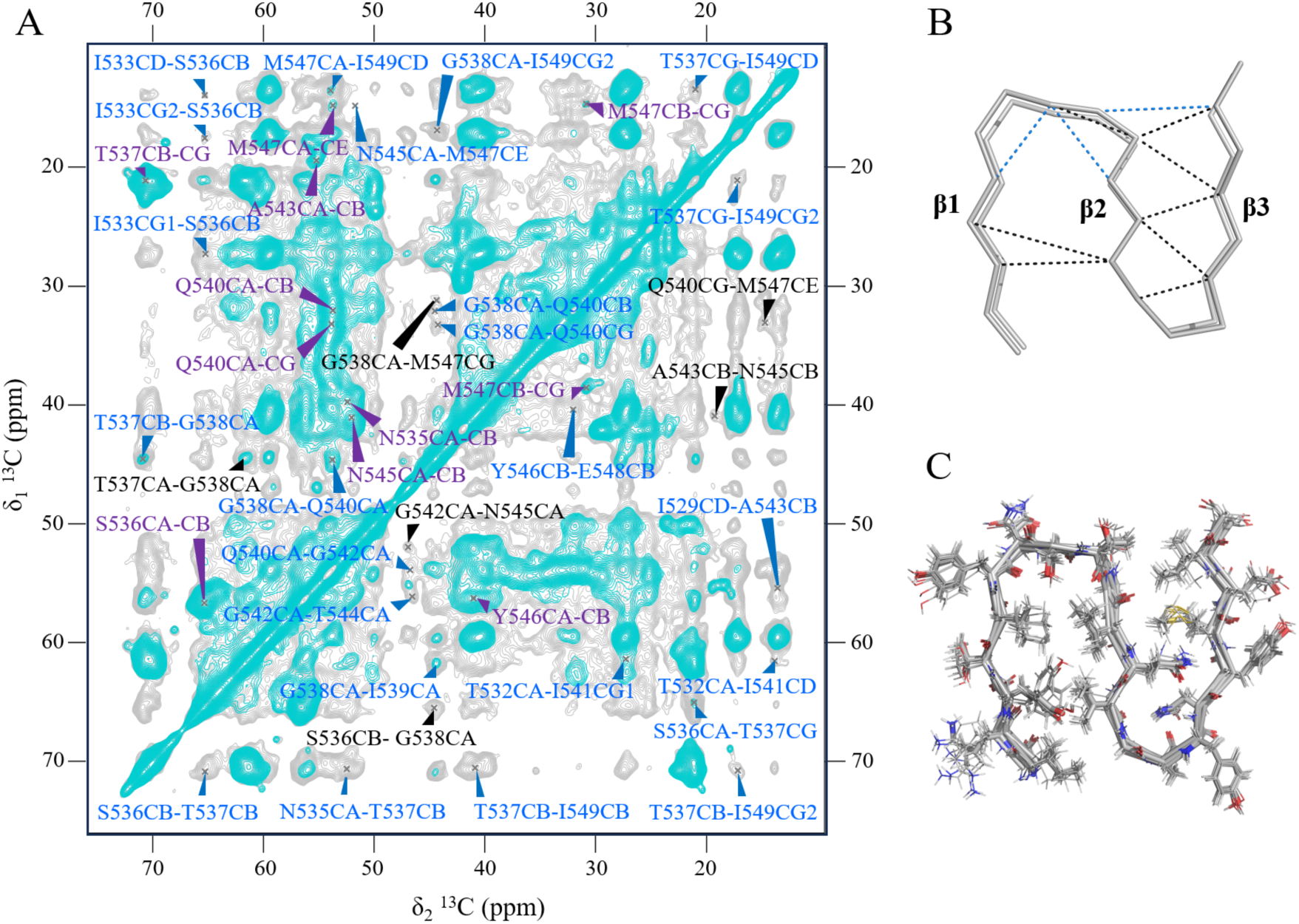
Structure calculation. **a** 2D ^13^C-^13^C CORD spectrum of hRIPK1 showing specific unambiguous (black), ambiguous (blue), and intraresidue cross peaks (purple) at 100 ms (cyan) and 500 ms (grey) mixing time. **b** Preliminary structure calculated using 36 torsion angle restraints and 7 unambiguous distance restraints (black dashed lines). Blue dashed lines represent low-ambiguity restraints that can be mapped onto the structure. **c** Full structure calculation with automatic assignment of all ambiguous distance restraints along with the restraints shown in panel b, and using this preliminary model as a starting structure or “seed”.

To construct the fibril model, long-range distance and torsion angle restraints were incorporated into structure calculations. Torsion angle restraints, phi and psi, were derived from TALOS+ predictions based on backbone chemical shifts, providing additional angular constraints that relate to the backbone fold. Long-range distance restraints were categorized as unambiguous or ambiguous, and their incorporation into the structural calculations was performed in two distinct stages. In the first stage, only unambiguous restraints were used to construct a preliminary structural model, ensuring that the core architecture of the amyloid fold was determined without the potential bias that ambiguous restraints could introduce. This initial model reveals that hRIPK1 fibril protomers display a three antiparallel extended β-strand architecture (**Fig. 3b**).

In a second stage, ambiguous restraints were incorporated to refine the model. These ambiguities were resolved through compatibility with the structural framework established in the first stage, yielding an ensemble of 10 structures that illustrate side chain orientations at the protomer level (**Fig. 3c**). This final model confirmed that hRIPK1 adopts an N-shaped fold within the amyloid core, spanning residues I529 to G550. Residues G551 and T552 were not included in the calculations as they lie outside of the fibril core and feature typical random coil chemical shift values (**Fig. 2b**).

The core comprises three antiparallel β-strands: β1 (I529-Y534, with a kink at N535-S536), β2 (T537-I541), which contains the RHIM motif I539-Q540-I541-G542 and with G542 located at a turn, and β3 (Y544-I549). The central β2 that harbors the RHIM motif serves as the primary stabilizing element, forming extensive interactions with β1 and β3. Specifically, β1 contributes residues Y531 and I533, which interact with residues I541 and I539 from β2 via hydrophobic contacts. A key hydrogen bond stabilizes the packing of β2 and β3, with Q540 mediating the interaction through its side chain and engaging the backbone carbonyl of Y546 (**Fig. 4**). The interface between β2 and β3 is further stabilized by the hydrophobic packing of the methylene groups of Q540 with the side chain from M547, while I549 encloses the fold, reinforcing this packing arrangement (**Fig. 4**).

**Fig. 4.**
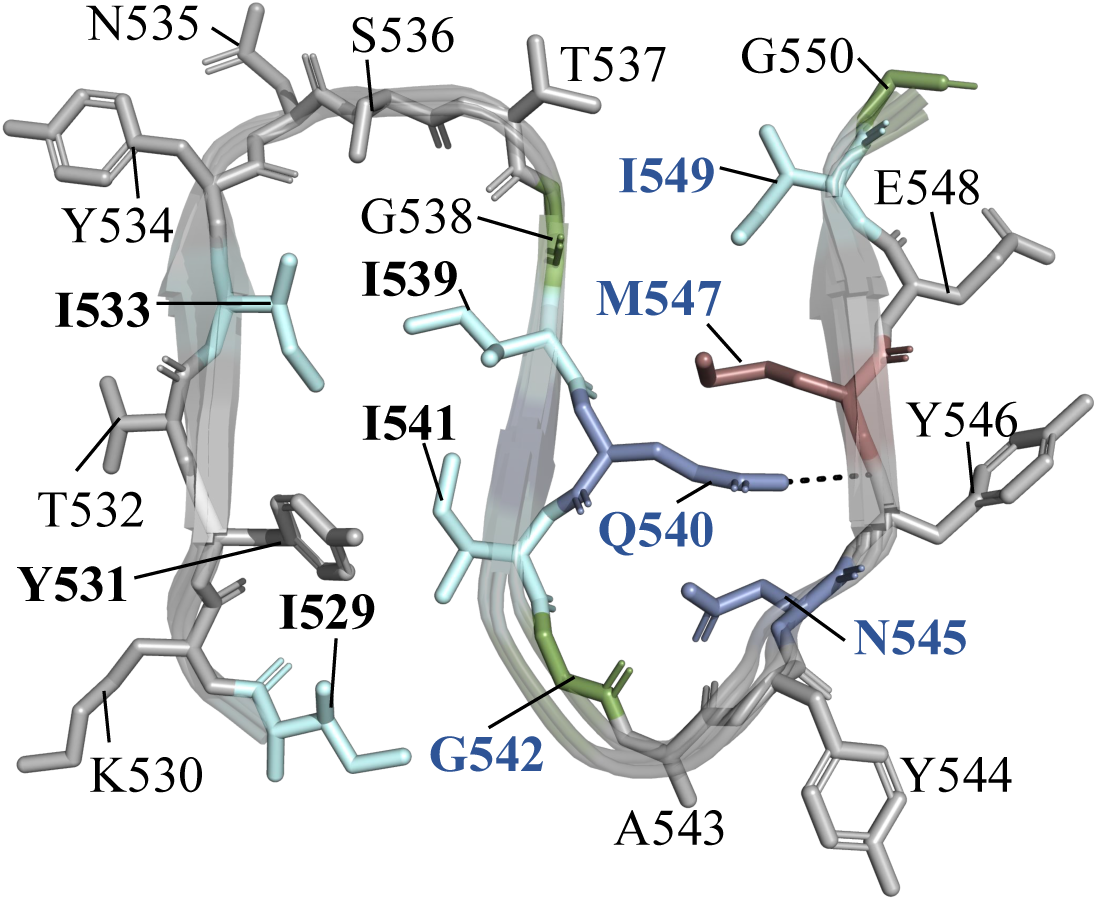
SSNMR structure of RHIM hRIPK1 protomer. Top view of the hRIPK1 (496–583) protomer derived from SSNMR, depicting the “N-shaped” fold formed by three β-strands. The central β-strand (β2) harbors the RHIM core tetrad IQIG. β1 and β2 enclose a hydrophobic core featuring four Ile (I529, I533, I539 and I541, in pale cyan) and a Tyr (Y531 in gray) residue, while β2 and β3 are linked together through an amide side chain to main chain H-bond (black dashed line) between Q540 (chromium) with Y546 (gray). The side chains of M547 (indium) and I549 (pale cyan) are also within the β2-β3 interface. Residues with black and blue bold labels correspond to the steric zippers between strands β1-β2 and β2-β3, respectively.

### Cryo-EM reveals the architecture of RHIM hRIPK1 fibrils

We employed cryo-EM to visualize how protomers organize to form RHIM hRIPK1 fibrils. Fibrils of hRIPK1 (496–583) dialyzed against water at a concentration of 80 µM were vitrified and imaged in-house using a Thermo Fisher Scientific 300 kV Krios G4 paired with a Gatan BioContinuum energy filter and K3 direct electron detector camera (see Materials and Methods). The 2.57 Å resolution cryo-EM density map captures hRIPK1 fibrils in a left-handed helical architecture with a width of 44 Å and a twist angle of -7.3° per layer and a rise of 4.67 Å that results in a pitch of 230.3 nm. The fibrils are assembled from protomers built by three antiparallel β-strands arranged in a N-shaped cross-β architecture (**Fig. 5a**). The high-resolution cryo-EM density map revealed well-resolved side-chain features for residues S528-G551, while the flanking regions could not be visualized presumably due to their flexibility in agreement with SSP values measured by SSNMR, and corroborated the presence of steric zippers of the RHIM motif-containing β2 strand with β1 and β3, as well as side chain-to-backbone hydrogen bonds connecting β2 and β3 that stabilize the amyloid core (**Fig. 5b**). Largely consistent with the fibrillar protomer structure determined by SSNMR (residues I529-G550), the structure visualized by cryo-EM provides additional information to understand protomer-protomer interactions and the assembly of RHIM hRIPK1 fibrils (**Fig. 5c-e**).

**Fig. 5.**
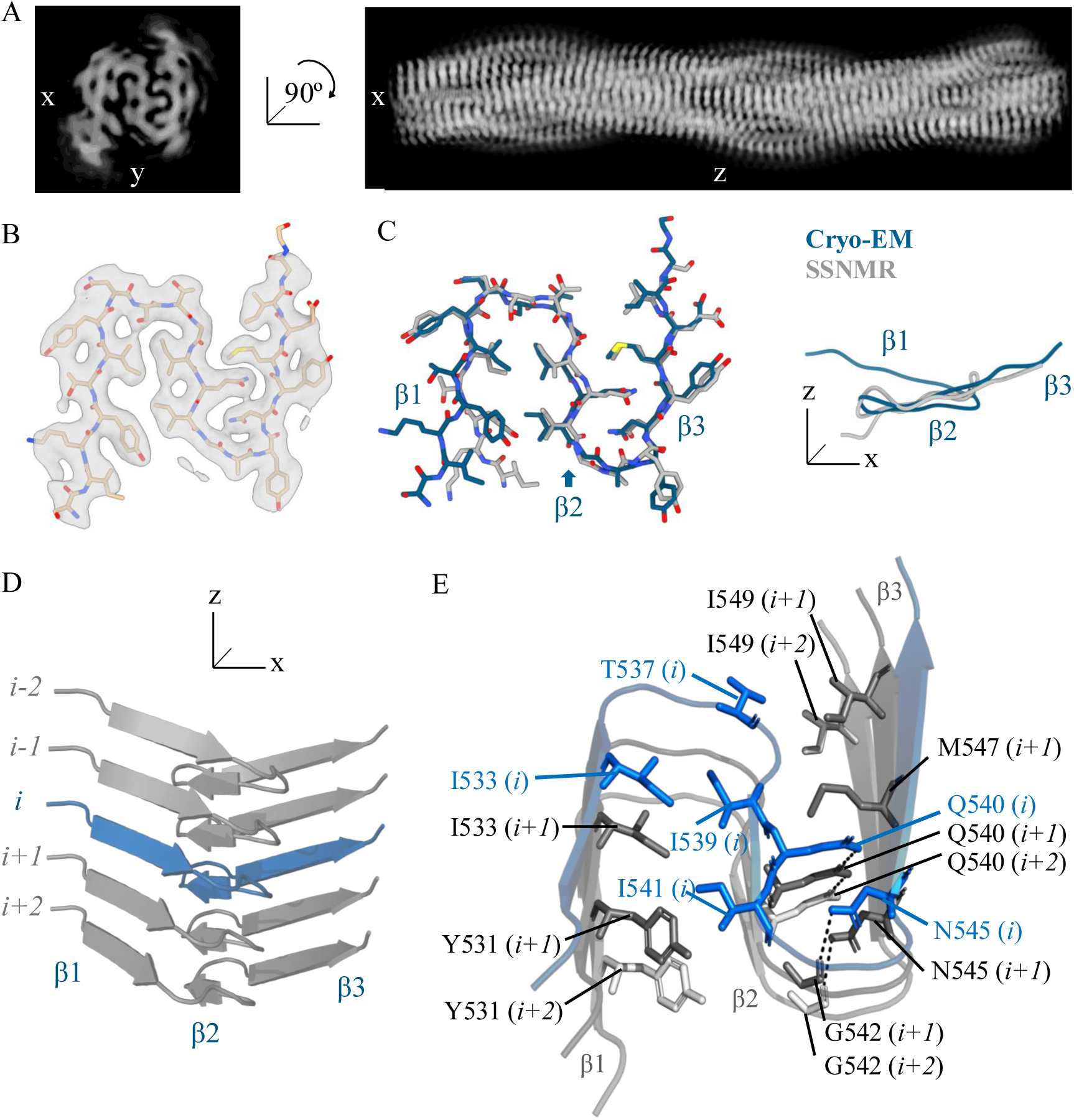
Cryo-EM structure of RHIM hRIPK1 fibrils. **a** Cross-sectional views (XY and XZ) of the 2.57 Å cryo-EM map of RHIM hRIPK1. **b** Top view of a single protomer of the density map of the fibril, with the atomic model of hRIPK1 fitted. **c** Structure alignment of both cryo-EM (blue) and SSNMR (grey) structures, viewed from different angles, highlighting the distinct planes of the three β-strands of a protomer in the cryo-EM structure. **d** Cartoon representation of five protomers (*i-2* to *i+2*), highlighting the overall fibril architecture. Protomer *i* is shown in blue, with adjacent protomers in grey. **e** Detailed view of interprotomer interactions between protomers *i* (blue), *i+1* and *i+2* (grey). Key interprotomer hydrogen bonds (e.g. N545-G542) are represented by black dashed lines, while hydrophobic contacts (e.g. Y531-I541, T537-I549) are illustrated by proximity between side chains.

The three antiparallel β-strands in each N-shaped hRIPK1 fibril protomer were found in distinct planes throughout the fibril axis, adopting a staggered arrangement that favors extensive interprotomer hydrogen bonding and hydrophobic contacts (**Fig. 5d**). Key interactions involving the RHIM-harboring β2 strand serve as the primary stabilizing element within the fibril, mediating contacts with β1 and β3 across protomers. At the β2-β1 interface, residue I541 in β2 of a protomer *i* establishes hydrophobic contacts with Y531 from adjacent protomers *i+1* and *i+2* (**Fig. 5e**). This staggered interaction anchors β2 to β1 of neighboring protomers, reinforcing the hydrophobic interface critical for fibril stability. Similarly, I539 in β2 of a same protomer *i* interacts with I533 from β1 of the same protomer, while also engaging I533 in the adjacent protomer *i+1*, further reinforcing the β1-β2 interface through overlapping stabilizing forces (**Fig. 5e**).

At the β2-β3 interface, Q540 in β2 of protomer *i* plays a dual role in fibril stabilization. First, it forms hydrophobic contacts with M547 from protomer *i+1*, locking β2 against β3 in adjacent protomers (**Fig. 5e**). Second, a ladder of Q540 side chain-to-side chain hydrogen bonds propagate along the fibril axis, linking protomer *i* to protomer *i+1* and continuing to *i+2*, creating a stabilizing zipper effect (**Fig. 5e**). In addition to Q540, the C-terminal end of β3 contributes to fibril closure. T537 in β2 of protomer i engages in hydrophobic interactions with I549 from protomers *i+1* and *i+2*, further linking β2 to β3 across protomers and effectively sealing the N-shaped fold.

A crucial hydrogen bonding network, centered around N545, underpins fibril integrity and facilitates elongation. N545 in β3 of protomer *i* forms a side chain-to-backbone hydrogen bond with G542 in β2 of protomer *i+1*, effectively locking β3 against β2 from adjacent protomers (**Fig. 5e**). This interaction propagates along the fibril, with each N545 engaging the G542 of the protomer directly below (e.g., N545 of *i+1* with G542 of *i+2*). Furthermore, N545 side chains stack along the fibril axis, contributing to the tight packing of adjacent layers. This stacking results in the side chain carbonyl group of N545 in protomer *i* forming a hydrogen bond with the amide backbone of N545 in protomer *i+1*, reinforcing interprotomer stability (**Fig. 5e**).

Together, these interprotomer interactions spanning hydrophobic packing and hydrogen bonding produce a tightly packed, highly stable fibril architecture that promotes elongation. This structural arrangement contrasts with hRIPK3 fibrils, where the three β-strands of a protomer lie on the same plane, and with N464 (equivalent to N545 in hRIPK1) adopting an outward-facing conformation that prevents hydrogen bonding with G461 (G542 in hRIPK1), which results in a more relaxed fibril architecture (**Fig. 6**).

**Fig. 6.**
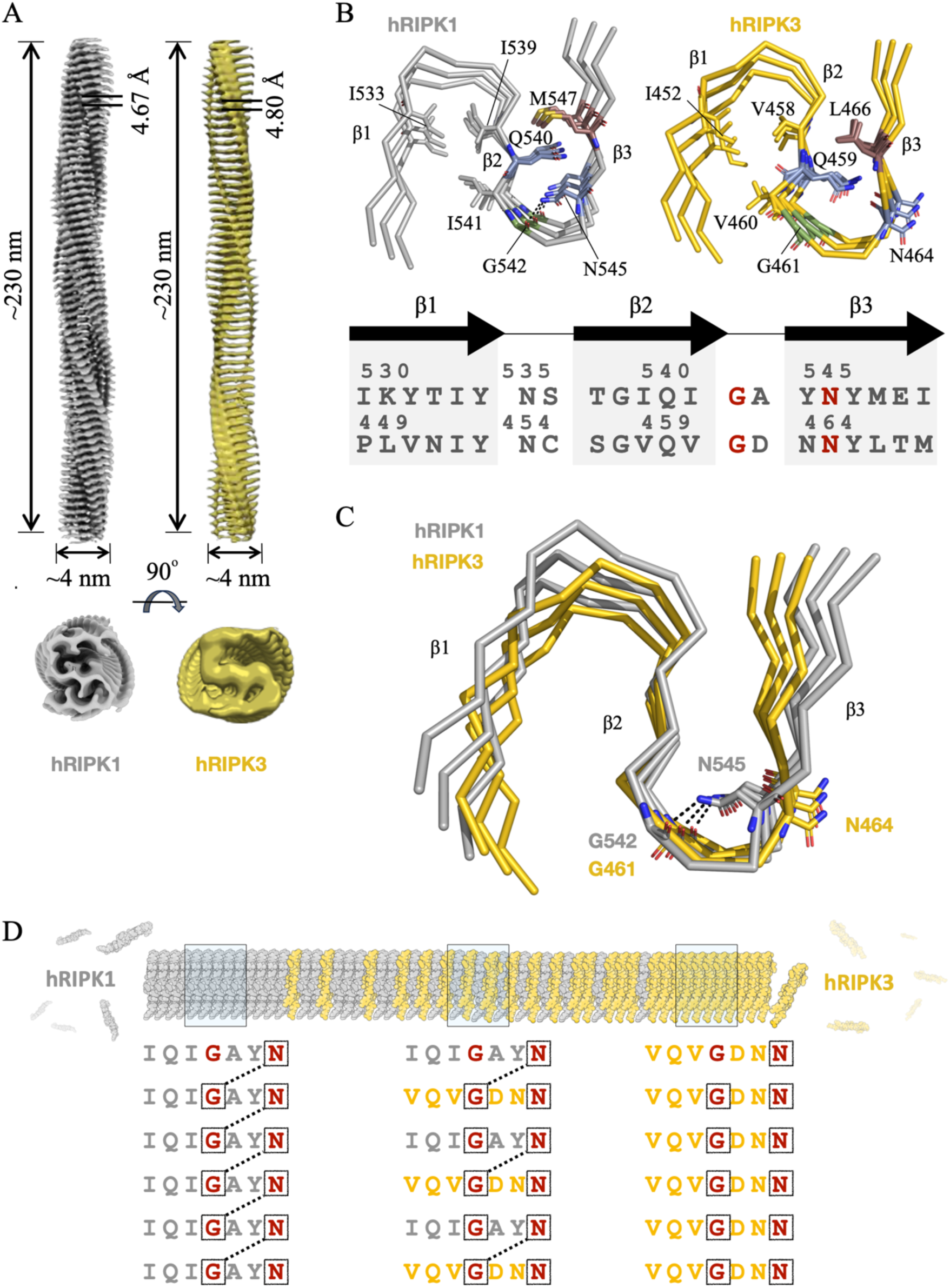
Proposed mechanism for hRIPK3 activation by hRIPK1. **a** Side and top views of the cryo-EM maps at 2.57 Å of hRIPK1 (left, grey) and at 4.24 Å of hRIPK3 (right, gold, adapted from ref. 22), showing pitch and width. **b** Cryo-EM structures of hRIPK1 (this work) and hRIPK3 adapted from ref. 22) alongside sequence alignment, highlighting the three β strands. In both structures, β strands 1 and 2 are stabilized by hydrophobic contacts between I533, I539 and I541 in hRIPK1, and their counterparts I452, V458 and V460 in hRIPK3. In hRIPK1, β strands 2 and 3 are stabilized by a side chain to backbone hydrogen bond mediated by Q540 (Q459 in hRIPK3), as well as hydrophobic packing between this residue and M547 (L466 in hRIPK3). N545 is buried within the steric zipper formed by β2 and β3, forming a side chain to backbone H-bond with G542, effectively closing the loop between β2 and β3. In contrast, the analogue residue in hRIPK3, N464, adopts an outward-facing orientation has a distinct orientation, preventing the formation of this hydrogen bond. **c** Detailed comparison of the N545-G542 hydrogen bond in hRIPK1 and the absence of this interaction between N464 and G461 in hRIPK3. **d** Proposed mechanism for hRIPK3 recruitment by hRIPK1, mediated by interprotomer hydrogen bonding between N545 (hRIPK1) and G461 (hRIPK3).

## Discussion

The advent of cold probes with unprecedented sensitivity for SSNMR has enabled faster and more cost-effective workflows for the structural characterization of biological systems (29). Here we employed CPMAS cryoprobe-detected SSNMR spectroscopy to map the amyloid core of RHIM hRIPK1 fibrils, identifying side-chain conformations that underpin the network of interactions stabilizing the compact structure of the fibril protomers. Cryo-EM revealed the architecture of RHIM hRIPK1 fibrils at high-resolution and the interactions between protomers underlying their assembly into helical fibrils. This integrative approach uncovered the structural basis for the architecture and assembly of amyloid fibrils formed by the master cell-signaling regulator hRIPK1.

hRIPK1 and hRIPK3 fibrils share a common structural framework of three antiparallel β-strands, with β2 harboring the RHIM motifs I539-Q540-I541-G542 in hRIPK1 and V458-Q459-V460-G461 in hRIPK3 (**Fig. 6a,b**). In both structures, β2 stabilizes β1 through hydrophobic residues (I533-I539-I541 in hRIPK1 and I452-V458-V460 in hRIPK3) and interfaces with β3 through Q540 and M547 in hRIPK1 (Q459 and L466 in hRIPK3) (**Fig. 6b**).

A defining difference between hRIPK1 and hRIPK3 fibrils lies in the interprotomer hydrogen bond between N545 (β3) and G542 (β2) in hRIPK1. This interaction is absent in hRIPK3, where N464 (analogous to N545) orients outward, preventing hydrogen bonding with G461 (analogous to G542) (**Fig. 6c**). This structural divergence results in the compact N-shaped fold observed in hRIPK1 fibrils, whereas the absence of the N464-G461 interaction in hRIPK3 yields a more relaxed S-shaped conformation. This greater conformational plasticity of hRIPK3 fibrils may facilitate the recruitment of MLKL or/and other downstream necroptosis effectors (14,15).

Despite these differences, the shared fibril architecture supports heteromeric assembly between hRIPK1 and hRIPK3. We propose that N545 in hRIPK1 initiates heteromeric fibril formation by engaging G461 in hRIPK3, effectively seeding its integration into the fibril core. This interaction is enabled by the outward-facing N464 in hRIPK3, which prevents hydrogen bonding with G461 across hRIPK3 protomers without altering the conserved backbone fold, thus allowing for staggered hRIPK1 protomer incorporation (**Fig. 6d**). As heteromeric fibrils elongate, hRIPK1 and hRIPK3 monomers may alternate along the fibril axis, following a 1:1 stoichiometry observed in prior studies (12). This staggered hydrogen bonding pattern could facilitate the transition from the compact N-shaped fold of hRIPK1 to the more open hRIPK3 conformation, echoing mechanisms described in other amyloid systems where minor side-chain differences govern cross-seeding specificity (31).

The critical role of N545 in hRIPK1 heteromeric fibril formation aligns with mutagenesis studies demonstrating that N545D disrupts hRIPK1-hRIPK3 assembly, while the analogous N464D mutation in hRIPK3 does not affect heteromeric fibril formation (11). This discrepancy reinforces the notion that N545 in hRIPK1 plays a pivotal role in initiating heteromeric fibril formation by engaging G461 in hRIPK3, while the outward-facing N464 in hRIPK3 does not contribute to the stabilization of such a heteromeric core.

The evolutionary conservation of Q540 and N545 in hRIPK1 – and their counterparts Q459 and N464 in hRIPK3 – underscores their importance in RHIM-mediated fibril formation. The distinct orientation of N464 in hRIPK3 may enhance fibril flexibility, facilitating MLKL recruitment. This flexibility likely explains why β3 residues of hRIPK3 were undetectable by CPMAS SSNMR in 1:1 hRIPK1:hRIPK3 heteromeric fibrils (12), whereas in hRIPK1 all residues in β3 remained well-defined. This structural adaptability may be key for transitioning from heteromeric to homomeric hRIPK3 fibrils, which are required for MLKL during necroptosis and other signaling pathways (17,18).

Although heteromeric amyloid assembly was first characterized in the functional RIPK1-RIPK3 complex, emerging evidence shows that this type of assembly may not be exclusive to physiological assemblies. Pathological amyloids involving TDP-43 and tau also form heteromeric assemblies in neurodegenerative diseases (32,33). Understanding how functional amyloids like hRIPK1 and hRIPK3 assemble and transition to heteromeric states will be essential to uncover which mechanisms may contribute to pathological aggregation via heterotypic recruitment into hybrid amyloids.

## Materials and Methods

### Protein Expression and Purification

The human RIPK1 RHIM domain (residues 496–583) was purchased from Genscript (New Jersey, NJ) with codons optimized for expression in *E. coli*, and subcloned in a pET11a derived vector include an N-terminal His×6 tag. This construct was cloned in BL21 (DE3). Samples uniformly labeled with isotopes (^13^C and ^15^N) were prepared following an adapted protocol described by Marley et al. (34) and Sivashanmugan et al. (35). In summary, transformed cells were cultivated in 2 L of LB medium until reaching an OD_600_ of 0.6–0.8. The cells were then harvested by centrifugation, and the resulting pellet was resuspended in 0.5 L of M9 minimal medium supplemented with ^13^C-glucose and ^15^NH_4_Cl (Cambridge Isotope Laboratories) as the sole nitrogen and carbon sources. To enhance isotope incorporation, the cells were incubated at 37 °C for 1.5 hours. Subsequently, the temperature was reduced to 25 °C for overnight protein induction using 0.5 mM IPTG.

For purification the cell pellet was resuspended in lysis buffer (50 mM Tris, 300 mM NaCl, and 1.3 µg/mL freshly prepared DNase) and lysed by sonication (30% amplitude, 5 seconds ON, 8 seconds OFF, for a total of 10 minutes) on ice. Cell debris was removed by centrifugation at 30,000 rpm for 20 minutes at 4°C, yielding an insoluble protein pellet (inclusion bodies). The inclusion bodies were resuspended and washed in a buffer containing 50 mM Tris, 1 mM freshly prepared DTT, and 1% Triton. To enhance resuspension, sonication was performed again (30% amplitude, 5 seconds ON, 8 seconds OFF, for a total of 10 minutes). The protein pellet was separated from the soluble fraction by centrifugation at 15,000 rpm for 15 minutes at room temperature. After this washing step, the remaining insoluble protein was treated with a buffer containing 1% SDS, 150 mM NaCl, 50 mM Tris (pH 8.0), and 1 mM freshly prepared DTT. The remaining insoluble material was resuspended in the same buffer (1% SDS, 150 mM NaCl, 50 mM Tris, pH 8.0, and 1 mM fresh DTT). The Ripk1 fusion protein was loaded onto a pre-equilibrated HisTrap column (GE Healthcare) to bind the His-tagged protein. A 5-column-volume (5 CV) washing step was performed using the resuspension buffer (1% SDS, 150 mM NaCl, 50 mM Tris, pH 8.0). A second washing step was carried out with 0.5% SDS, 150 mM NaCl, and 50 mM Tris (pH 8.0). Elution was performed using the same buffer supplemented with 0.5 M imidazole (0.5 M imidazole, 0.5% SDS, 150 mM NaCl, 50 mM Tris, pH 8.0).

### Fibril Assembly

For fibril preparation, two different protocols were followed depending on the intended application. For SSNMR experiments, ^13^C, ^15^N isotopically labeled and unlabeled proteins were mixed at a 1:4 ratio in 50 mM Tris-HCl (pH 7.4), 150 mM NaCl, and 2% SDS to achieve the desired isotopic dilution while maintaining the proteins in non-assembled states. For cryo-EM fibrils, unlabeled protein was kept in in 50 mM Tris-HCl (pH 7.4), 150 mM NaCl, and 8M urea, also ensuring the proteins remained unassembled. In both cases, the protein mixtures were subsequently dialyzed against Milli-Q water (SSNMR samples) or 50 mM Tris-HCl (cryo-EM samples), at pH 7.4, using a 3500 Da molecular weight cutoff dialysis membrane (Spectrum^TM^ 123110). The water being replaced every 24 hours at room temperature to remove the denaturant and allow fibril assembly.

### Thioflavin T Fluorescence Assay

The ThT fluorescence assay was performed following the procedure of LeVine (27). A 1.0 mM stock solution of ThT (Sigma, St. Louis, MO) in water (M.Q.) and pH 7.4 was prepared. Samples for fluorescence measurements were prepared by mixing 5 μL of the ThT stock solution with protein samples, resulting in a final ThT concentration of 50 μM. The protein concentration was adjusted to 25 μM in a final volume of 200 μL. Fluorescence measurements were conducted at 25 °C using a Jobin-Yvon Fluoromax-4 spectrofluorometer. The excitation wavelength was set to 440 nm, and emission spectra were recorded from 460 to 500 nm at a scan speed of 2 nm·s⁻¹. Both excitation and emission slit widths were set to 3 nm.

### Circular Dichroism Spectroscopy

Far-UV CD spectra were recorded using a JASCO J-710 spectropolarimeter. Protein samples were prepared at a concentration of 50 μM in water (M.Q.), pH 7.4. Measurements were performed with a bandwidth of 1.2 nm and a scan speed of 20 nm·min⁻¹. The spectra were collected over a wavelength range of 190–260 nm at 25 °C, using a quartz cuvette with a path length of 0.1 cm. Each spectrum was obtained by averaging eight scans to minimize noise. A spectrum of the buffer was recorded under identical conditions and subtracted from the sample spectra to eliminate background contributions.

### Negative-Stain Transmission Electron Microscopy (TEM)

Human RIPK1 fibrils were characterized by negative-stain TEM to evaluate their morphology. A 5 μL aliquot of the fibril preparation was applied on a 400-mesh copper grid coated with a carbon-plated supporting film and negatively stained by incubation with 2% uranyl acetate for 2 minutes before blotting off to remove excess stain. Grids were previously treated with a glow discharge protocol of 20 mA during 30 s to achieve negative polarity in their surface and promote fibril deposit. After three washes with deionized water and blotting the excess water, the grid was left to air dry for 5 minutes at room temperature.

TEM analysis was performed using a TALOS L12C transmission electron microscope equipped with a Ceta-F 16M digital camera. Images were acquired at a magnification of 28,000x and an accelerating voltage of 120 kV.

### Solid-State NMR Spectroscopy

Dialyzed fibrils from hRIPK1 were pelleted and transferred into a Bruker 3.2-mm rotor using Bruker packing tools and a Ortoalresa Minicen RT255 centrifuge for SSNMR analyses. Excess water was removed during the packing process. Approximately, 60 mg of wet fibrils were packed (ca. 15 mg of labelled material).

SSNMR experiments were conducted using a Bruker AVANCE NEO spectrometer operating at a magnetic field strength of 14.1 T (600 MHz ^1^H frequency) and a HCN CPMAS cryogenically-cooled probe for high sensitivity ^13^C detection (29). Three 3D spectra were recorded to determine sequential connectivity, namely NCACX, NCOCX, and CANCOCX experiments with 50 ms CORD mixing (36–38). To assess longer-range correlations, 2D CORD spectra were acquired with mixing times of 50 ms, 100 ms, and 500 ms. The long mixing time (500 ms) provided critical distance constraints between β2 with β1 and β3 that define the monomer fold in its fibrillar conformation. All experiments were collected at physiologically relevant temperatures (ca. 37°C), and the spectra were indirectly referenced to DSS. Spectra were processed using Topspin 4.0 (Bruker Biospin) and analyzed with Sparky (D. Goddard and D. G. Kneller, SPARKY 3, University of California, San Francisco). Complete experimental details and acquisition parameters are provided in **Table S1**.

### Structure Calculation

The core structure of the hRIPK1 amyloid fibril, encompassing residues I529 to I549, was calculated using CYANA 3.97. Two rounds of calculations were performed to achieve the final atomic-resolution structure.

The initial calculation began with two non-interacting, extended hRIPK1 molecules subjected to simulated annealing calculations using 20000 torsion angle dynamics steps to censure proper sampling of the conformational space. During this calculation, 100 structures were generated using the following restraints: 36 dihedral angle restraints derived from TALOS+ predictions (39) based on backbone chemical shifts and 7 unambiguous restraints with a cutoff of 8.0 Å. Since the overall monomeric fibrillar conformations of the backbone in amyloids are already well-defined with a limited number of unambiguous cross-peaks (40), a preliminary structure calculation already defined the N-shaped amyloid core shown in **Fig. 3b**. From this preliminary structural model, 11 low-ambiguity restraints could be resolved by visual inspection of compatibility (**Table S2**). Low-ambiguity restraints correspond to cross-peaks where the assignment in one dimension is unambiguous while there are two possible assignments in the other dimension (i.e. two possibilities for that cross-peak). A test calculation with these 14 restraints gave confidence on the preliminary model. In a second stage, a full CYANA calculation with seven cycles of combined automatic cross-peak assignment and structure calculation was performed using a tolerance of 0.2 ppm for matching input cross-peak positions with the chemical shifts of potential assignments, while keeping the cutoff of 8.0 Å for distance restraints with a calibration search from 3 to 10 Å. A total of 100 structures were generated, from which the 10 lowest-energy models were selected as the final representative ensemble. Both stages of structure calculation incorporated a symmetric dimeric model, in order to restrain the protomeric chain lying within a same plane. To this end, hydrogen bonding patterns predicted from secondary structure data were imposed as additional restraints. Specifically, backbone CO and N upper limits of 3.2 Å were imposed between residues exhibiting negative SSP values, characteristic of β-sheet alignment, following the protocol by Schütz et al. (40). These restraints ensure the fidelity of parallel β-sheets and maintain the monomers in a planar configuration and aligned consistently within the fibril, mimicking the periodicity of hydrogen bonds in fibrils.

A detailed summary of the structural statistics, including restraint violations, RMSD values, and refinement parameters, is provided in **Table S3**. The validated structural ensemble supports the “N-shaped” amyloid fold of hRIPK1 fibrils, consistent with independent cryo-EM reconstructions.

### Cryo-EM Imaging and Data Processing

Samples for cryo-EM were vitrified in a FEI Vitrobot Mark IV with 100% humidity at 4°C by applying 4 μL hRIPK1 fibril solution (final concentration 80 μM) to mesh 300 R1.2/1.3 gold grids (UltrAuFoil). The grids were freshly glow-discharged before sample application for 3 min at 15 mA and 4.2 × 10^−1^ mbar in an ELMO glow discharge system (Cordouan Technologies, France). Grids were blotted for 5.5 s using a blotting force of 2.

Automated data acquisition was carried out on a 300 kV Krios G4 electron microscope (Thermo Fisher Scientific) equipped with a BioContinuum/K3 direct detector (Gatan) operating in counting mode at a calibrated pixel size of 0.5054 Å. A -0.8 to -1.6 μm defocus range was used to automatically record 8,650 movies of 50 frames using EPU 2 (Thermo Fisher Scientific) with aberration n-free image shift and fringe-free imaging. The electron dose rate was set to 14.2 e−/pixel/s, with a total exposure time of 0.88 s, resulting in a cumulative dose of 49.3 e−/Å² per micrograph. Initial frame alignment and contrast transfer function (CTF) estimation was performed using CryoSPARC Live (41). The best movies according to CTF and astigmatism were chosen for particle selection. Particle picking was performed in cryoSPARC (41) with the filament tracer tool using templates generated from manual picking and a distance between segments of 10 Å. A total of 2,814,672 filament segments were picked. Several rounds of reference-free 2D class averaging were used to clean the dataset to a final number of 460,173 segments (**Fig. S1**).

### Helical Reconstruction

An initial model was built using *ab initio* reconstruction in Cryosparc that was used as a template for a homogeneous refinement without imposing helical symmetry. A symmetry search job was carried out to determine the starting helical parameters (a helical rise of 4.09 Å and a twist angle of -6.53°) that were refined in subsequent non-uniform refinements. To further improve the model, several rounds of local and global CTF refinement (42), as well as reference-based motion correction were applied. The final map has a helical rise of 4.667Å and a twist angle of -7.319°, and a resolution of 2.57 Å based on the gold-standard Fourier shell correlation (FSC) = 0.143 criterion. The resulting map revealed detailed molecular features, consistent with the cross-β architecture observed in hRIPK1 fibrils.

### Model Building

An initial atomic model was generated using ModelAngelo (43) with the 2.57 Å resolution map and the sequence of the protein. All fibril chains except the five central ones were removed from the initial model. The retained chains were subjected to several iterative cycles of real-space refinement in Phenix (44), followed by manual adjustments to the map using Coot (45) to improve the fit and optimize the geometry. Once the central five-chain model was refined, all chains except the central one were removed. Symmetry operations were then applied using Phenix to reconstruct the complete fibril. The cryo-EM figures were prepared using ChimeraX (46), and the single layer of the density map was extracted using the Segger tool (47).

### Validation and Deposition of SSNMR and Cryo-EM structures

The resulting structures were validated using the RCSB PDB validation server (https://validate-rcsb-2.wwpdb.org/). The final 10 SSNMR models were deposited in the Protein Data Bank (PDB) under accession code 9HR9. The corresponding NMR chemical shifts were deposited in the Biological Magnetic Resonance Data Bank (BMRB) under accession number 34971. Cryo-EM atomic coordinates have been deposited at the Protein Data Bank (PDB) and cryo-EM maps at Electron Microscopy Data Bank with accession codes 9HR6 and EMD-52356, respectively.

## Acknowledgements

This work is funded by the European Union (ERC, 101042403 – BiFOLDOME, to M.M). Views and opinions expressed are however those of the authors only and do not necessarily reflect those of the European Union or the European Research Council. Neither the European Union nor the granting authority can be held responsible for them. NMR experiments were performed in the “Manuel Rico” NMR Laboratory (LMR) of the Spanish National Research Council (CSIC), a node of the Spanish Large-Scale National Facility (ICTS R-LRB). G.T.A.-D. and P.P. acknowledge FJC2021-047976-I fellowships funded by MCIN/AEI/ 10.13039/501100011033 and European Union Next Generation EU/PRTR, and PIPF-2022/BIO-25611 funded by the Dept. of Education, Science and Universities from the Community of Madrid, respectively. This work has been supported by grant PID2022-143177NB-I00 to I. U.-B from the Spanish Ministry of Science, Innovation and Universities. Cryo-EM was performed at the Basque Resource for Electron Microscopy located at Instituto Biofisika (UPV/EHU, CSIC), supported by the Department of Science, Universities and Innovation and the Innovation Fund of the Basque Government, with additional support from MCIN (Recovery, Transformation and Resilience Plan) and the Basque Government “Biotechnology Complementary Plan Applied to Health” with funding from European Union NextGenerationEU (PRTR-C17.I1; PRTR-C17.I01.P01.S13) (AAAA_ACG_AY_2539/22_05). We thank also Igor Tascón (Instituto Biofisika) for interesting discussions on this work.

## Author Contributions

P.P., F.C.E.-G., and G.A.T.-D prepared the samples for SSNMR, CD and ThT, and conducted these experiments. P.P. assigned all the SSNMR spectra and prepared fibrils amenable for cryo-EM. M.M. performed SSNMR structure calculations and wrote the original draft. J.P.L.-A., I.U.-B., H.J., conducted cryo-EM sample preparation, data collection, analysis and structure determination. Funding acquisition and project management: M.M. and I.U.-B. P.P., J.P.L.-A., I.U.-B., H.J., F.C.E.-G., G.A.T.-D., and M.M, prepared the figures and edited the final draft.

## Conflict of Interest

The authors declare no conflict of interest.

## Notes

### Competing Interest Statement

The authors have declared no competing interest.

