## Supplementary Information for "Structural Basis for the Assembly of Amyloid Fibrils by the Master Cell-Signaling Regulator Human Receptor-Interacting Protein Kinase 1"

**Figure S1 | Workflow of cryo-EM data processing and helical reconstruction**. A total of 2,814,672 filament segments were automatically picked from 8,650 micrographs and subjected to iterative 2D classification to remove noise and poor-quality segments. An initial 3D model was generated using ab initio reconstruction without imposing symmetry constraints. Helical parameters (rise and twist) were determined through a symmetry search, followed by refinement using non-uniform reconstruction. To improve the map, several rounds of local and global CTF refinement, as well as reference-based motion correction, were performed. The local resolution of the final map, estimated using the 0.5 FSC criterion, is shown.

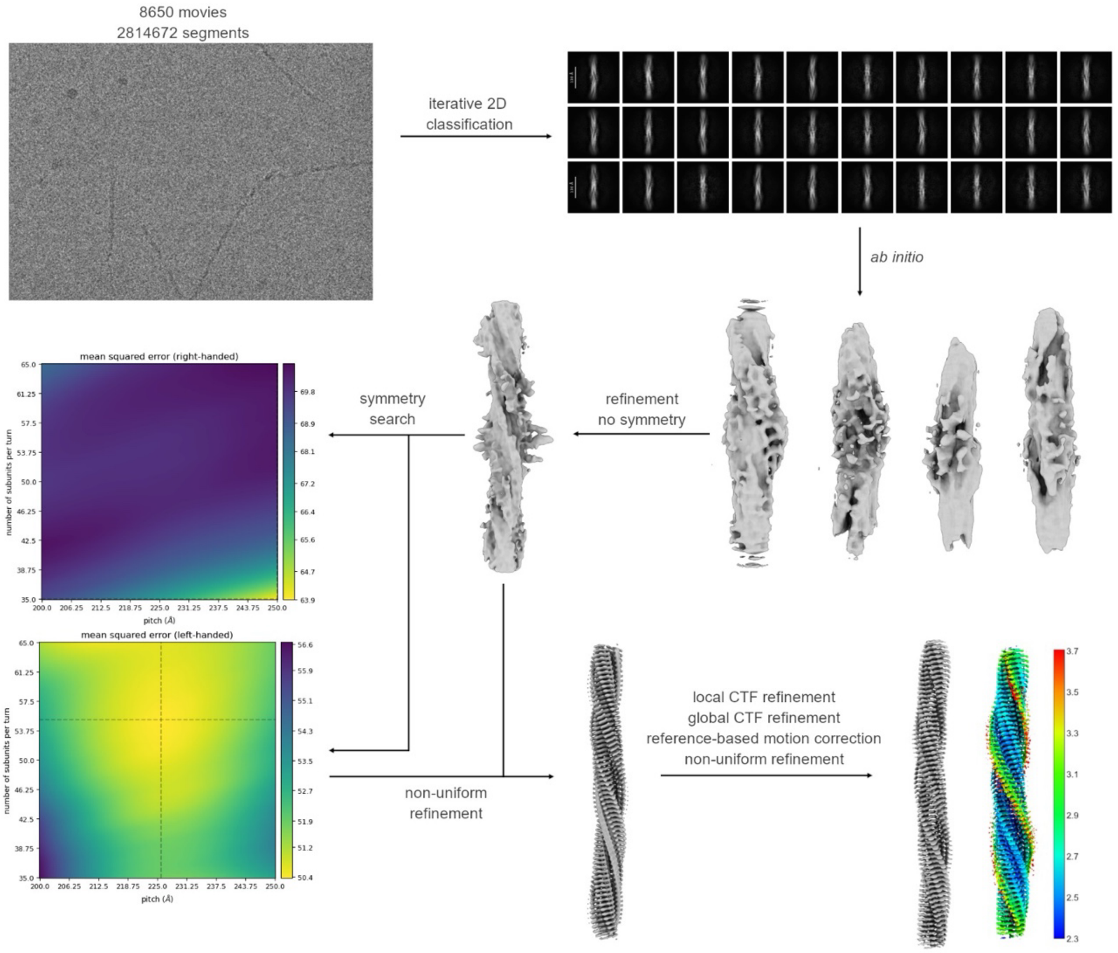

**Table S1** | SSNMR acquisition parameters.

| 14.1 T (600 MHz 1H frequency), 3.2 mm HCN CPMAS CryoProbe | | | | | | | | | | |
|---|---|---|---|---|---|---|---|---|---|---|
| Experiment | NS | MAS (kHz) | d1 (s) | Acq time (ms) | Sweep width (kHz) | Dec power (kHz) | Mixing (ms) | CP time (ms) | Field Strength (kHz) | Sample T (K) |
| 2D NCA | 8 | 14 | 2 | t2: 15; t1: 11 | ω2: 52.6; ω1: 2.8 | 100 | - | HN: 1.00; NC: 1.50 | H: 54; N: 40 (transfer 1); N: 4; CA: 18 (transfer 2) | 315 |
| 2D NCACX | 8 | 14 | 2 | t3: 15.0; t1: 7.9 | ω3: 52.6; ω2: 3.5; ω1: 2.8 | 100 | 50 | HN: 1.00; NC: 1.50 | H: 54; N: 40 (transfer 1); N: 4; CA: 18 (transfer 2) | 315 |
| 3D NCACX | 8 | 14 | 2 | t3: 15.0; t2: 8.0; t1: 7.9 | ω3: 52.6; ω2: 3.5; ω1: 2.8 | 100 | 50 | HN: 1.00; NC: 1.50 | H: 54; N: 40 (transfer 1); N: 4; CA: 18 (transfer 2) | 315 |
| 3D NCOCX | 8 | 14 | 2 | t3: 15.0; t2: 8.0; t1: 7.9 | ω3: 52.6; ω2: 3.5; ω1: 2.8 | 100 | 50 | HN: 1.00; NC: 1.50 | H: 54; N: 40 (transfer 1); N: 8.4; CA: 5.6 (transfer 2) | 315 |
| 3D CANCOC | 64 | 14 | 2 | t3: 17.4; t2: 7.9; t1: 6.0 | ω3: 58.8; ω2: 2.8; ω1: 4.7 | 100 | 50 | HC: 1.20; CN: 1.25; NC: 7.00 | H: 69; CA: 55 (transfer 1); CA: 3; N: 11 (transfer 2); N: 8.4; CO: 5.6 (transfer 3) | 315 |
| 2D CORD | 4 | 14 | 2 | t2: 20; t1: 17 | ω2: 44.2; ω1: 35 | 100 | 5, 20, 100, 500 | HC: 1.75 | H: 69 ; C: 55 | 315 |

**Table S2** | Restraints identified in the spectra.

**Unambiguous restraints**

| Residue 1 | Residue 1 type | Atom 1 type | Residue 2 | Residue 2 type | Atom 2 type | |
|---|---|---|---|---|---|---|
| 532 | THR | C | 541 | ILE | CD | * |
| 536 | SER | CA | 538 | GLY | CA | * |
| 536 | SER | CB | 538 | GLY | CA | * |
| 538 | GLY | CA | 550 | GLY | CA | * |
| 538 | GLY | CA | 547 | MET | CG | * |
| 538 | GLY | CA | 547 | MET | CE | * |
| 538 | GLY | C | 547 | MET | CE | * |
| 540 | GLN | CG | 545 | ASN | CB | * |
| 540 | GLN | CG | 547 | MET | CE | * |
| 540 | GLN | CD | 547 | MET | CE | * |
| 542 | GLY | CA | 545 | ASN | CA | * |
| 543 | ALA | CB | 545 | ASN | CB | |

**Ambiguous restraints**

| Residue 1 | Residue 1 type | Atom 1 type | Residue 2 | Residue 2 type | Atom 2 type | |
|---|---|---|---|---|---|---|
| 533 | ILE | CA | 536 | SER | CB | |
| 533 | ILE | CB | 536 | SER | CB | |
| 533 | ILE | CG1 | 536 | SER | CB | ** |
| 533 | ILE | CG2 | 536 | SER | CB | ** |
| 533 | ILE | CD | 536 | SER | CB | ** |
| 533 | ILE | CD | 536 | SER | CA | ** |
| 535 | ASN | CA | 537 | THR | CB | |
| 536 | SER | CA | 539 | ILE | CG1 | ** |
| 536 | SER | CB | 539 | ILE | CA | |
| 536 | SER | CB | 539 | ILE | CB | |
| 536 | SER | CB | 539 | ILE | CG1 | ** |
| 536 | SER | CB | 539 | ILE | CG2 | ** |
| 537 | THR | CA | 535 | ASN | CA | |
| 537 | THR | CB | 535 | ASN | CA | |
| 537 | THR | CA | 539 | ILE | CG1 | |
| 537 | THR | CB | 539 | ILE | CG1 | |
| 537 | THR | CA | 539 | ILE | CD | |
| 537 | THR | CB | 539 | ILE | CD | |
| 537 | THR | CG | 539 | ILE | CD | |
| 537 | THR | CB | 550 | GLY | CA | |
| 537 | THR | CG | 550 | GLY | CA | ** |
| 537 | THR | CB | 549 | ILE | CB | |
| 537 | THR | CB | 549 | ILE | CG2 | |
| 537 | THR | CG | 549 | ILE | CG2 | |

| | | | | | | |
|---|---|---|---|---|---|---|
| 537 | THR | CG | 550 | ILE | CD | |
| 538 | GLY | CA | 540 | GLN | CA | |
| 538 | GLY | CA | 540 | GLN | CB | |
| 538 | GLY | CA | 540 | GLN | CG | |
| 538 | GLY | CA | 547 | MET | CA | |
| 538 | GLY | CA | 547 | MET | CB | |
| 538 | GLY | CA | 549 | ILE | CG2 | |
| 540 | GLN | CA | 542 | GLY | CA | |
| 540 | GLN | CA | 547 | MET | CE | ** |
| 540 | GLN | CB | 547 | MET | CE | ** |
| 542 | GLY | CA | 544 | TYR | CA | |
| 542 | GLY | CA | 545 | ASN | CB | ** |
| 545 | ASN | CA | 547 | MET | CE | |
| 546 | TYR | CB | 548 | GLU | CB | |
| 547 | MET | CA | 549 | ILE | CD | |

In light green: restraints between nuclei that are 2 residues apart in the sequence

In dark green: restraints between residues that are 3 or more residues apart in the sequence

* Distance restraints used in first structure calculation

** Low ambiguity restraints between nuclei that are 3 or more residues apart and that can be resolved based on the preliminary model

**Table S3 |** SSNMR structure calculation statistics.

<table>
<tr><th>Statistic description</th><th>Value</th></tr>
<tr><td>Total distance restraints</td><td>121</td></tr>
<tr><td>Intra-residue (|i-j|=0)</td><td>4</td></tr>
<tr><td>Sequential (|i-j|=1)</td><td>44</td></tr>
<tr><td>Medium range (|i-j|>1 and |i-j|<5)</td><td>48</td></tr>
<tr><td>Long range (|i-j≥5)</td><td>25</td></tr>
<tr><td>Total dihedral-angle restraints</td><td>35</td></tr>
<tr><td>Number of restraints per residue</td><td>7.1</td></tr>
<tr><td>Number of long range restraints per residue</td><td>1.1</td></tr>
</table>

**Table S4** | Cryo-EM data collection, refinement and validation statistics

| | Human RIPK1 (EMDB-52356) (PDB 9HR6) |
|---|---|
| **Data collection and processing** | |
| Magnification | 165000 |
| Voltage (kV) | 300 |
| Electron exposure (e–/Å$^2$) | 49.3 |
| Defocus range (μm) | 0.8-1.6 |
| Pixel size (Å) | 0.5054 |
| Axial symmetry | C1 |
| Twist (degree) | -7.319 |
| Rise (Å) | 4.667 |
| Initial segment images (no.) | 2814672 |
| Final segment images (no.) | 460173 |
| Map resolution (Å) | 2.57 |
| FSC threshold | 0.143 |
| Map resolution range (Å) | 2.32-2.86 |
| **Refinement** | |
| Initial model used | *In silico* (ModelAngelo) |
| Model resolution (Å) | 2.57 |
| FSC threshold | 0.143 |
| Model resolution range (Å) | 2.32-2.86 |
| Map sharpening *B* factor (Å$^2$) | -57.4 |
| Model composition | |
| Non-hydrogen atoms | 930 |
| Protein residues | 120 |
| Ligands | 0 |
| *B* factor (Å$^2$) | 65.70 |
| R.M.S. deviations | |
| Bond lengths (Å) | 0.004 |
| Bond angles (°) | 0.789 |
| Validation | |
| MolProbity score | 0.82 |
| Clashscore | 0 |
| Poor rotamers (%) | 0 |
| Ramachandran plot | |
| Favored (%) | 95.45 |
| Allowed (%) | 4.55 |
| Disallowed (%) | 0.00 |
